# Integration of Machine Learning Improves the Prediction Accuracy of Molecular Modelling for *M. jannaschii* Tyrosyl-tRNA Synthetase Substrate Specificity

**DOI:** 10.1101/2020.06.26.174524

**Authors:** Bingya Duan, Yingfei Sun

## Abstract

Design of enzyme binding pocket to accommodate substrates with different chemical structure is a great challenge. Traditionally, thousands even millions of mutants have to be screened in wet-lab experiment to find a ligand-specific mutant and large amount of time and resources is consumed. To accelerate the screening process, here we propose a novel workflow through integration of molecular modeling and data-driven machine learning method to generate mutant libraries with high enrichment ratio for recognition of specific substrate. *M. jannaschii* tyrosyl-tRNA synthetase (*Mj.* TyrRS) is used as an example system to give a proof of concept since the sequence and structure of many unnatural amino acid specific *Mj.* TyrRS mutants have been reported. Based on the crystal structures of different *Mj.* TyrRS mutants and Rosetta modeling result, we find D158G/P is the critical residue which influences the backbone disruption of helix with residue 158-163. Our results show that compared with random mutation, Rosetta modeling and score function calculation can elevate the enrichment ratio of desired mutants by 2-fold in a test library having 687 mutants, while after calibration by machine learning model trained using known data of *Mj.* TyrRS mutants and ligand, the enrichment ratio can be elevated by 11-fold. This molecular modeling and machine learning-integrated workflow is anticipated to significantly benefit to the *Mj.* tyrRS mutant screening and substantially reduce the time and cost of web-lab experiment. Besides, this novel process will have broad application in the field of computational protein design.

**CCS Concepts:** • Applied computing • Life and medical sciences • Computational biology • Molecular structural biology

## 1 Introduction

Genetic code expansion technology^1^ has been widely used in biology research which can be applied on monitoring protein conformational change^2^ caused by PTM^3^ as biophysical probe^4^, improving enzyme activity^5^ and designing proteins with novel catalytic functions^6–8^. Through this technology, we can incorporate artificially designed unnatural amino acid (UAA) into almost any specific site of target protein. Usually an orthogonal tRNA and amino acyl-tRNA synthetase (aaRS) mutant pair is necessary for recognition of specific UAA and subsequently acyl ligation to tRNA. Though there are more than 100 UAAs and their corresponding orthogonal aaRS mutants have been reported, UAAs with novel chemical structure, biophysical and biological function will significantly benefit to the biology research and protein therapeutic^9^. Pursue of the UAA’s corresponding aaRS by researchers has never stopped. Traditionally, an aaRS mutant recognizing specific UAA is selected by several rounds of positive and negative screening from large and diverse aaRS mutant libraries rationally designed from wild type aaRS^1^. The screening process is very tedious, manpower-costing and error-prone. As a result, a more rapid and efficient method to identify aaRS mutant for specific UAA is in urgent need.

From the perspective of computational chemistry, finding aaRS mutant for specific UAA is a protein design problem. The sequence space of wild type aaRS as receptor is explored with the goal of finding the mutant having lowest binding free energy to the UAA ligand. In recent years, computational chemistry based-molecular modelling method, such as protein homology modelling^10^, protein structure prediction^11^, molecular docking^12^ and protein design^13^ has been rapidly developed. Large number of successful cases on enzyme design for substrate selectivity^14^ are reported.

Recently there have been great advances on artificial intelligence (AI) technology, such as machine learning and deep learning. AI has been used in computer vision, speech recognition, machine translation, small-molecule drug design^15^, protein engineering^16^, antibody structure prediction^17^ and antibody design^18^. In general AI can extract representative features of protein sequence and structure from large amount of chemical molecule and protein data, learn internal patterns which can’t explicitly spotted by human and utilize experimental validated properties such as binding affinity data, IC50 and enzyme activity as labels to train a model which can be used to predict which sample has target property of interest from huge number of molecules and diversified protein mutant libraries. For example, improved score function of molecular docking software^19^, improved protein design performance^20^ and accurate prediction of thermostability for protein mutants^21^ have been achieved through machine learning and deep learning. Combined with computational chemistry-based method, data-driven AI method is giving satisfactory solution and accurate prediction on many biological problems.

In this work, we collected all the *M. jannaschii* tyrosyl-tRNA synthetase (*Mj.* TyrRS) mutants reported in the literature to compare and analyze the sequence and structural feature and difference between mutants and wild type *Mj.* TyrRS. Then we use Rosetta EnzymeDesign method^22^ to model the structure and predict binding pose of UAA-*Mj.* TyrRS mutant pair. Finall, machine learning model is integrated to calibrate the score function for aaRS substrate selectivity prediction and mutant design to achieve better prediction accuracy. We think the improved *Mj.* TyrRS-UAA selectivity prediction model will be extremely useful for *in silico* screening of UAA-specific aaRS mutants.

## 2 Results

### 2.1 Crystal structures comparison between *Mj.* TyrRS mutants and WT

Firstly, we collected the information of all UAA and corresponding *Mj.* TyrRS mutants of which high-resolution X-ray crystal structure has been released, as shown in Scheme 1, Table 1 and Supplementary file S1. In total there are 52 UAAs and 132 TyrRS mutants. The X-ray crystal structure of 18 TyrRS mutants has been reported (Table 1). The diversity of UAAs (Scheme 1) is large. Both of small and large, polar and non-polar, hydrophilic and hydrophobic substitution groups of p-hydroxy of Tyr are included in these UAAs. In the WT of *Mj.* TyrRS, hydrogen bond network between tyrosine hydroxy group and pocket residues stabilizes the tyrosine ligand and lowers the binding free energy (Figure 1). In order to recognize other UAAs in Scheme 1, mutations with different amino acid types and physicochemical properties have to be introduced to accommodate different chemical properties.

**Table 1.**
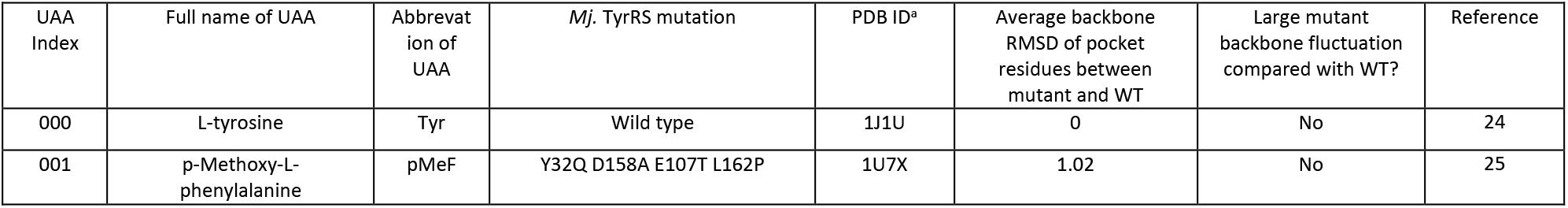
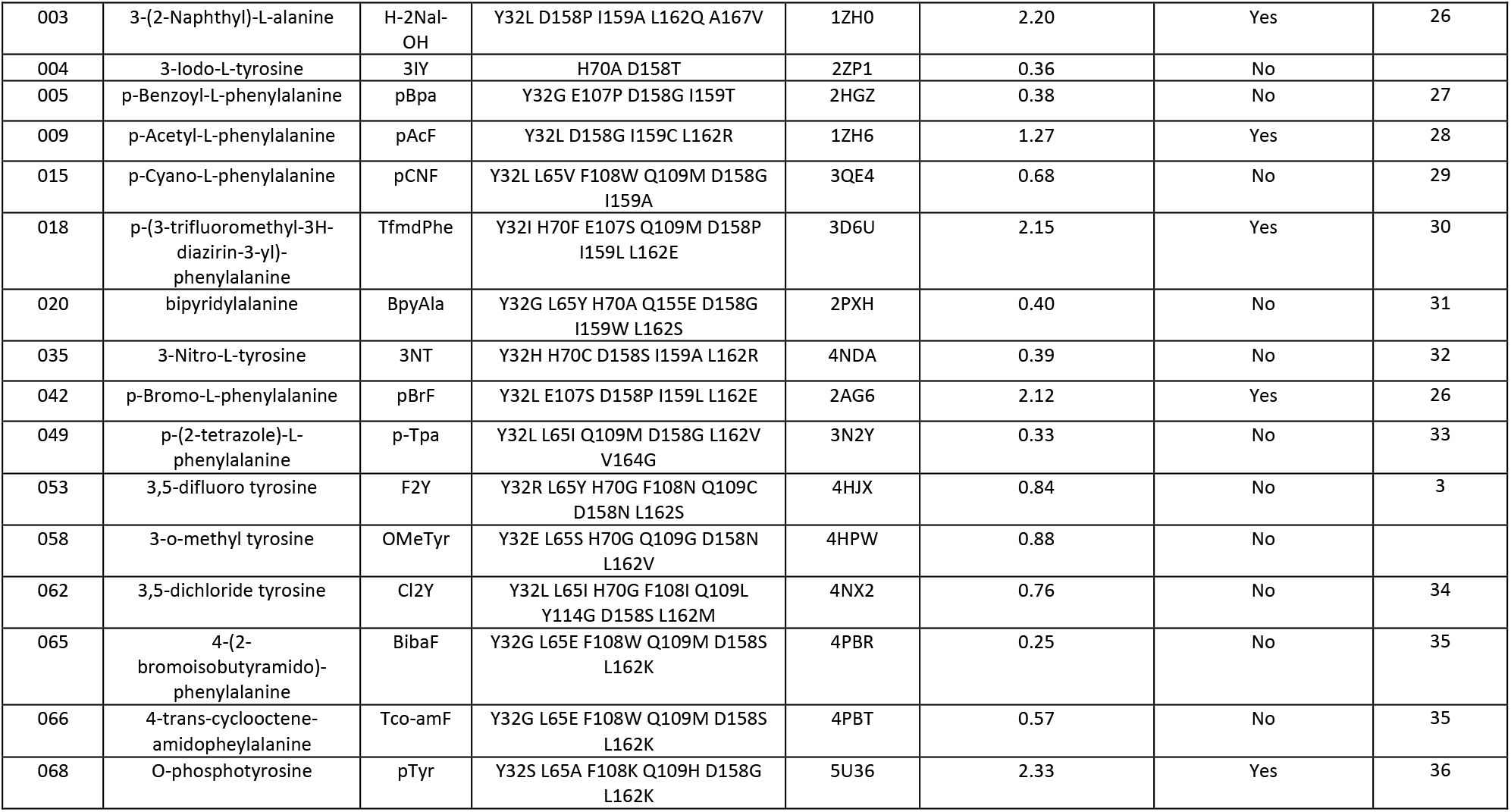
UAA index and name, aaRS mutation, PDB ID and X-ray crystal structure backbone RMSD of UAA binding pocket residues. Crystal structures of mutants with the same sequence is only shown once.

**Scheme 1.**
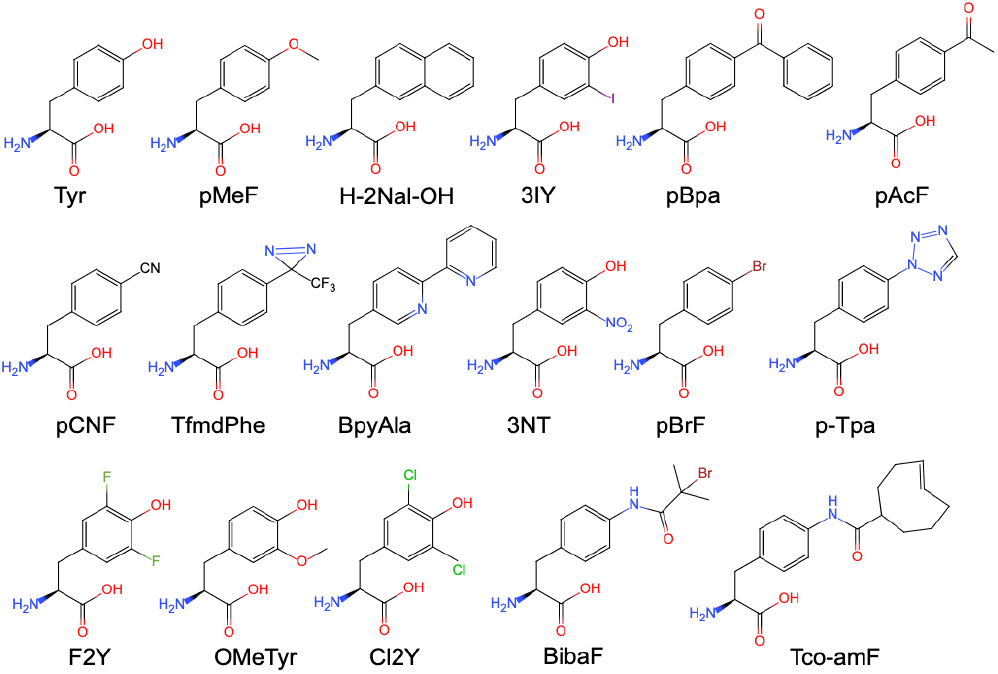
Chemical structure of the UAAs recognized by *Mj.* TyrRS mutants of which X-ray crystal structure has been solved.

**Figure 1.**
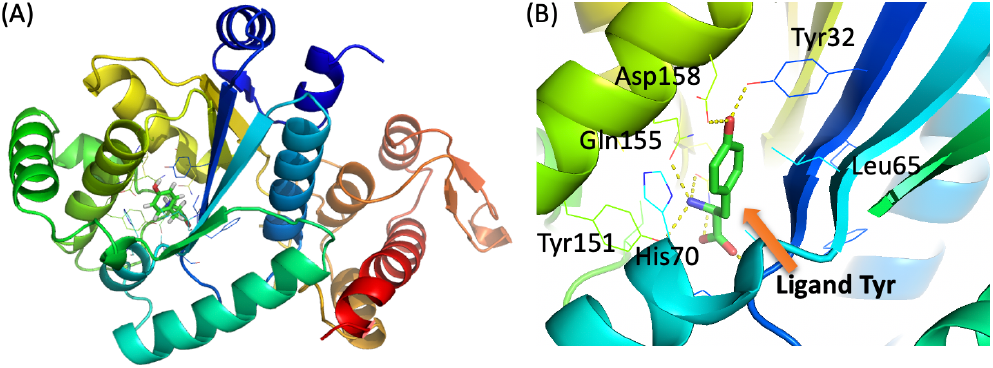
Crystal structure of the WT *Mj.* TyrRS and tyrosine complex. PDB ID: 1J1U. The figure is rendered by Pymol^23^.(A) *Mj.* TyrRS is shown as cartoon. Tyrosine ligand is shown as sticks. Surrounding residues are shown as lines. (B) The binding pocket of ligand tyrosine. Tyrosine forms hydrogen bond network with Tyr32, Asp158 and Gln155, which stabilizes the hydroxy group of tyrosine.

For a reliable model of the mutant structure, the backbone fluctuation introduced by mutation should be small compared with that of WT. It has been reported that some mutations can change the backbone conformation^36^, so next we aligned the crystal structures of *Mj.* TyrRS mutants and WT and then calculated the RMSD of backbone N, CA, C, O of binding pocket residues between mutant and WT to compare the structural difference. As shown in Table 1 and Figure 2A, structures with PDB ID 1ZHO, 1ZH6, 2AG6, 3D6U, 3D6V have average backbone RMSD larger than 1.2, of which backbone conformation is changed by mutation compared with WT. Other mutants have small backbone fluctuation which can be ignored when building the homology model. 4 of the 16 mutant structures have large backbone fluctuation (Table 1). In Figure 2B, backbone RMSD of different binding pocket residues are compared. We can see residues 158~163 have larger RMSD than other residues, which are on the alpha helix near UAA ligand. 4 representative mutant structures are chosen and superimposed on WT structure, as shown in Figure 3. In the structure of 1ZH0 and 2AG6 the backbone change of alpha helix is large while in 2HGZ and 4PBR the change is small. This implies it may be difficult to obtain accurate UAA and pocket residue binding pose prediction result for mutant bearing large backbone conformation change in homology model.

**Figure 2.**
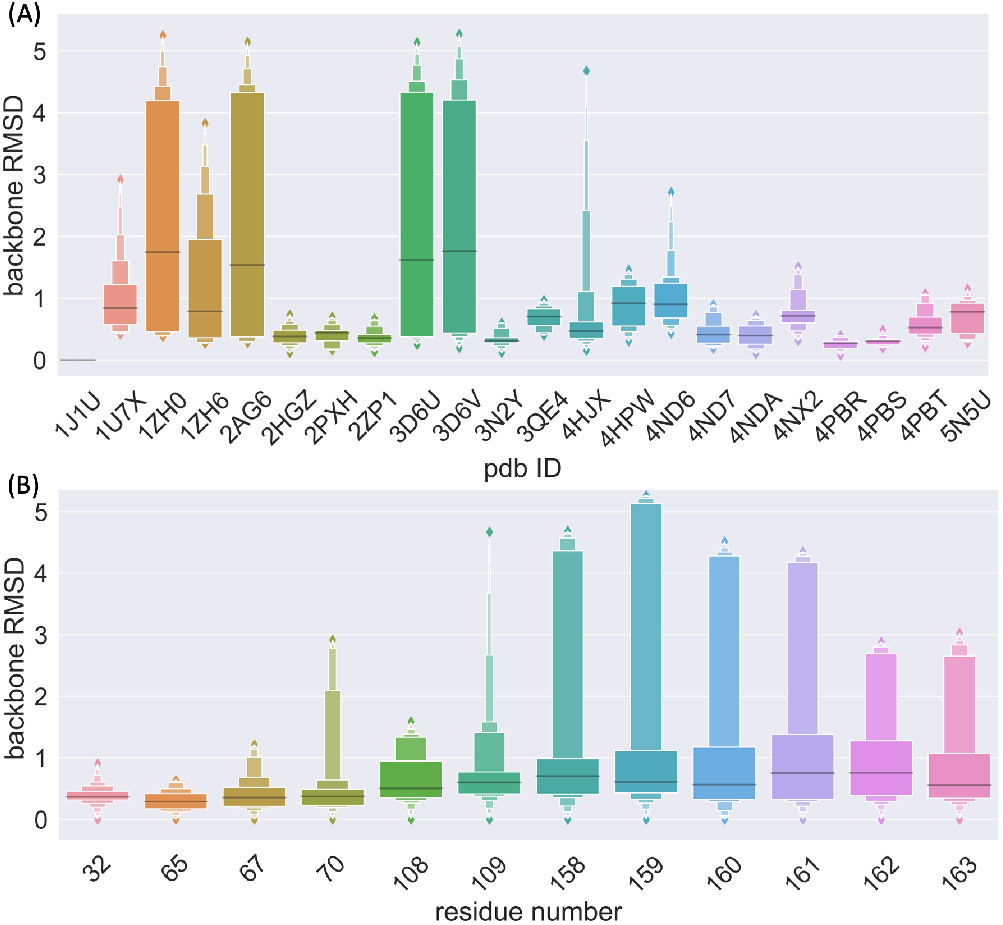
Crystal structure backbone heavy atom RMSD of UAA binding pocket residues (12 in total) between mutants (21 in total) and WT. (A) For each mutant, backbone RMSD of pocket residues is calculated and plotted using boxen plot showing different quantiles. (B) For each pocket residue, backbone RMSD from all mutants and WT is calculated and shown.

**Figure 3.**
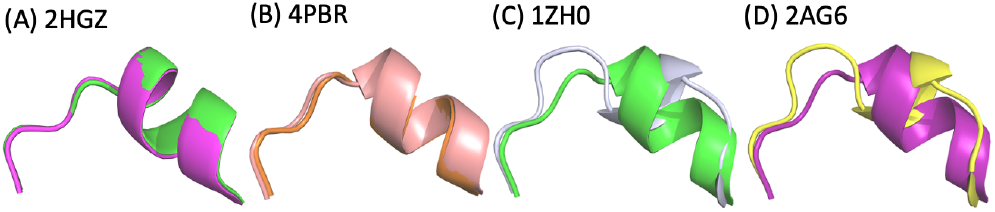
Comparison of the alpha helix containing residues 155-166 between 4 representative *Mj.* TyrRS mutants and WT X-ray crystal structures. WT structure is colored in magenta, light pink, green, magenta for (A), (B), (C) and (D), respectively.

How different mutations induce the backbone conformation change remains unclear. It was reported that in pTyr *Mj.* TyrRS D158G mutation acted as a helix breaker and caused the rearrange of the helix and opened up the pocket to accommodate the bulky UAA^36^. Though pCNF *Mj.* TyrRS has the same D158G mutation, no obvious backbone conformation change is observed (Table 1). To further investigate the critical residues affects the helix conformation, we calculated AAindex ^37^(531 types of numerical indices representing various physicochemical and biochemical properties of amino acids) of residues 158~163 for the 17 *Mj.* TyrRS in Table 1. Totally there are 531 * 6=3186 features for each TyrRS mutant. We performed analysis of variance (ANOVA) by sklearn^38^ to find which residue and AAindex property will contribute most to the discrimination of TyrRS helix backbone disruption. The top largest 10 f_value are from residue 158 and have p_value < 0.001. The 10 AA index types are related to intrinsic secondary structure propensities of the amino acids (Table 2), which implies that residue 158 may influences the secondary structure. AAindex type and value versus backbone disruption or not for 17 pdbs and residue 158 is shown in Figure 4. We can see that the distribution of the AAindex value is different for TyrRS with different helix backbone. Each of the 10 AAindex can be used to separate the TyrRS mutant with helix backbone disruption from others. This indicates that D158P/G mutation is a helix breaker and can disrupt the backbone conformation, which should be taken into consideration when building homology model for the TyrRS mutants to obtain accurate structure model.

**Table 2.** Residue site and top 10 AAindex contributes most to the discrimination of TyrRS helix backbone disruption.

| Position | AAindex type | $f_{\text{value}}$ | $p_{\text{value}}$ |
| --- | --- | --- | --- |
| 158 | MUNV940104 | 27.74547 | 0.000063 |
| 158 | ROBB760104 | 27.262113 | 0.000069 |
| 158 | MUNV940105 | 26.026208 | 0.000089 |
| 158 | BLAM930101 | 25.948507 | 0.00009 |
| 158 | BUNA790103 | 25.905193 | 0.000091 |
| 158 | QIAN880134 | 25.607837 | 0.000097 |
| 158 | MUNV940101 | 24.323819 | 0.000126 |
| 158 | ONEK900101 | 23.726363 | 0.000144 |
| 158 | RACS820114 | 23.355244 | 0.000156 |
| 158 | QIAN880112 | 22.67982 | 0.000181 |

**Figure 4.**
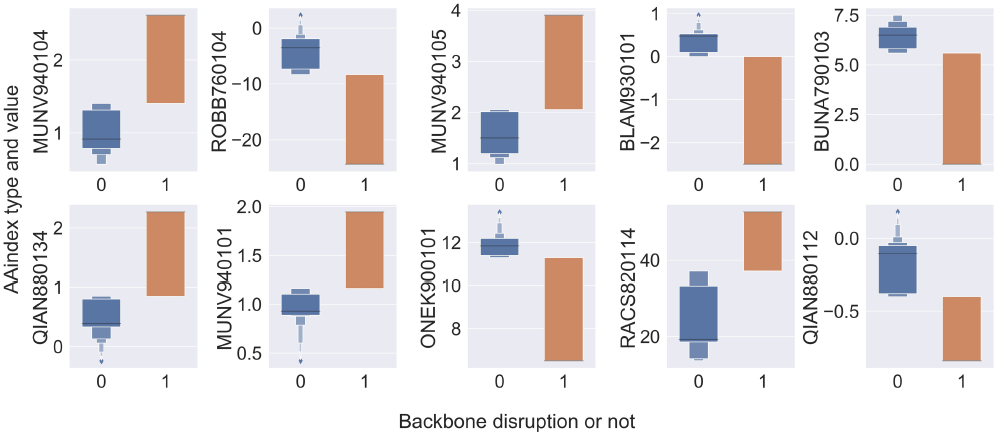
Boxplot of AAindex type and value versus backbone disruption or not for 17 pdbs and residue 158. Each subplot has a different AAindex type. There are 17 data points in each plot, which is the AAindex value of residue 158 from 17 pdbs. 1 is for backbone disruption while 0 is not.

### 2.2 UAA-Mj. TyrRS mutant complex modelling using Rosetta

To facilitate the TyrRS wet-lab screening process and reduce the time and resource cost, molecular modelling is carried out to predict which TyrRS mutant can recognize the target UAA. First we use Rosetta EnzymeDesign^22^ to model the backbone change and AA side chain packing to accommodate specific UAA ligand for each of the 6864 UAA-mutant pair complex (52 UAAs times 132 TyrRS mutants). The crystal structure of WT *Mj.* TyrRS (PDB ID 1J1U) is used as input template. Representative modeling results are shown in Figure 5. For mutants with PDB ID 2HGZ and 4PBR, no obvious backbone disruption is observed in crystal structure (Table 1). The predicted binding pose in Figure 5A 2HGZ is accurate, while in Figure 5B 4PBR the predicted orientation of UAA ligand deviates from the true position in crystal structure and leads to inaccurate side chain packing of residue Lys162 and Ser158, which may be caused by wrong selection of BibaF conformation. For mutants with PDB ID 1ZH0 and 2AG6, large backbone disruption is observed in crystal structure (Table 1). Though the UAA ligand position and conformation (Figure 5C and 5D) is accurately modeled, the side chain of AA surrounding the binding pocket deviates a lot from the crystal structure mainly due to inaccurate modeling of the alpha helix with residues 158-163. The results indicate that for mutants with little disruption on helix 158-163, Rosetta model is accurate enough to predict the binding pose but for mutants with large disruption, the opposite is true.

**Figure 5.**
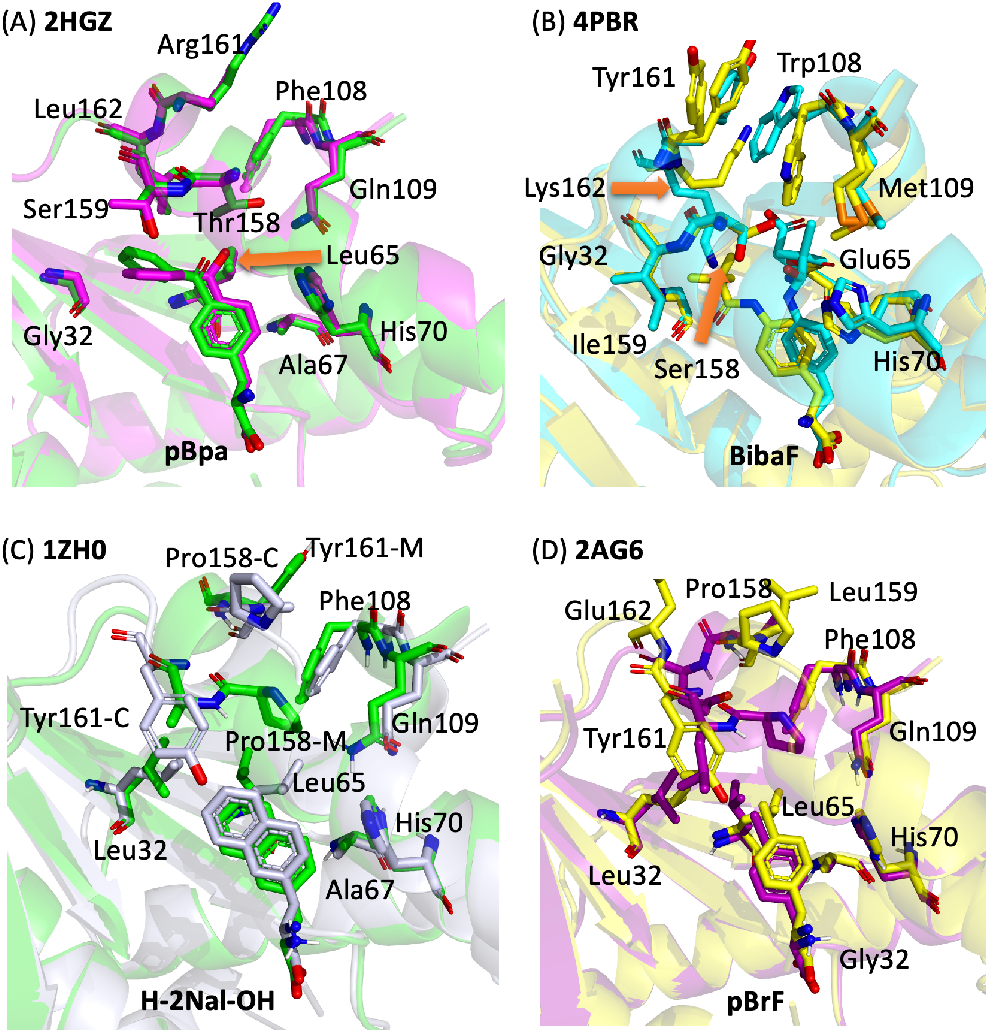
Comparison of the binding pose and binding pocket residues of 4 *Mj.* aaRS mutants from X-ray crystal structure and Rosetta model. Corresponding UAA and binding site residues are shown as sticks. (A) PDB ID: 2HGZ. Crystal: green. Model: magenta. (B) PDB OD: 4PBR. Crystal: orange. Model: light pink. (C) PDB ID: 1ZH0. Crystal: white. Model: green. (D) PDB ID: 2AG6. Crystal: yellow. Model: magenta.

To test whether the correct backbone conformation of residues 158-163 helix having D158G/P mutation can be predicted, this segment is *de novo* remodeled using KIC loop modeling method^39^ in Rosetta. 1000 models are generated for each of the 4 mutants with PDB ID 1ZH0, 1ZH6, 2AG6 and 3D6U using homology model in previous step as input. The backbone RMSD to crystal structure are calculated and plotted against total energy (Figure 6). Scatter plot in funnel shape can be observed for all 4 mutants, which indicates the amount of backbone conformation sampling is enough to find local minimum energy point. For the lowest energy structure, the RMSD to crystal structure is 3.65, 2.41, 3.74 and 4.39 Å in the 4 mutants. The lowest RMSD to crystal structure in 1000 models of the 4 mutants is 0.94, 0.47, 0.49 and 0.48, respectively. The rmsd_to_crystal for the top 10 models ranked by total_energy_score and total_score rank for the top 10 models ranked by rmsd_to_crystal on 4 TyrRS mutants are shown in Figure 7A and 7B, respectively. To further analyze the modeled structure, crystal structure, input Rosetta model, model with lowest RMSD to crystal, model with lowest energy are superimposed. We can see that model for 1ZH6 has the most ideal funnel plot (Figure 6B) and model with the lowest energy has least deviation from crystal structure (Figure 7D), though 2.41 Å RMSD may still not be accurate enough for UAA-TyrRS binding affinity prediction. For the remaining 3 mutants, model structure close to the native crystal structure has been generated but we can’t pick it up due to the error of Rosetta energy function (Figure 7).

**Figure 6.**
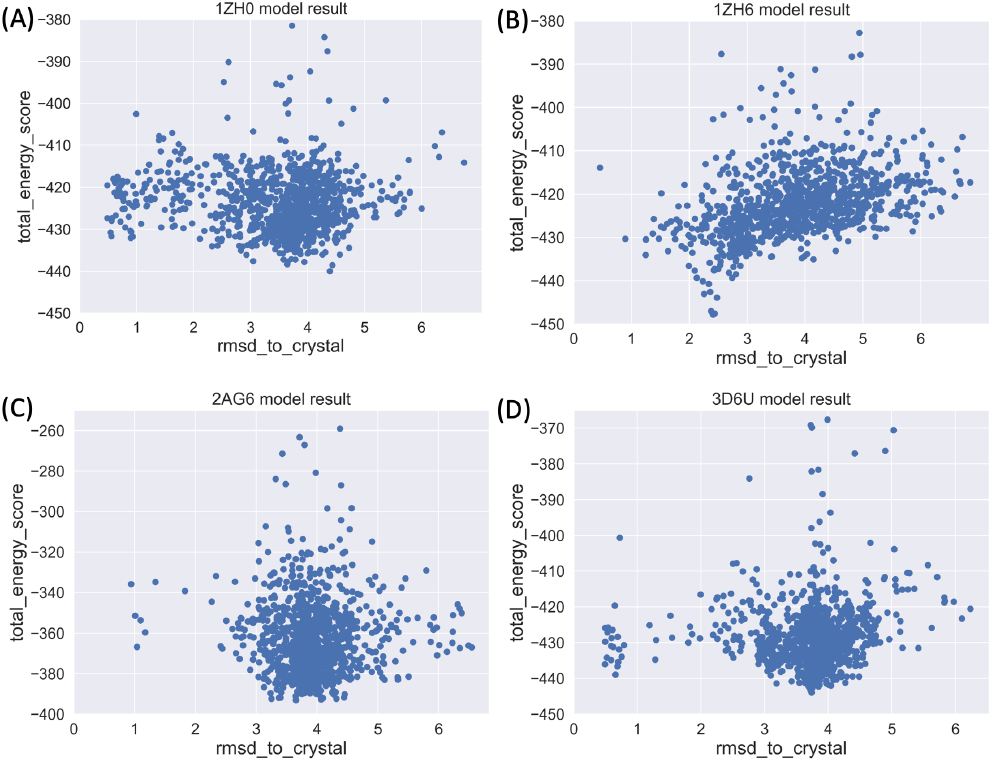
Scatter plot for rmsd_to_crystal vs total_energy_score in 1000 models generated by Rosetta *de novo* loop modelling for TyrRS mutants with PDB ID 1ZH0, 1ZH6, 2AG6 and 3D6U.

**Figure 7.**
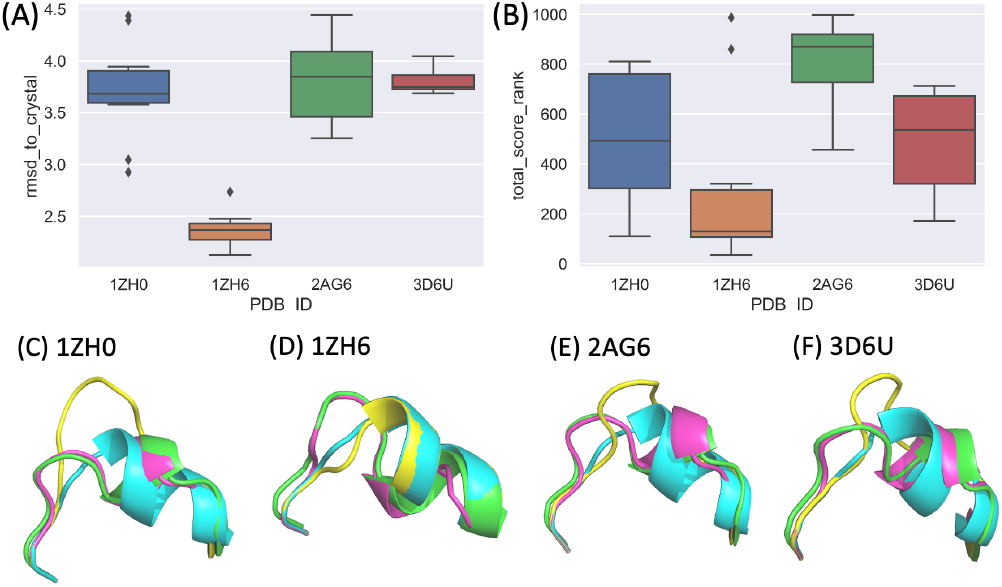
Structure analysis of Rosetta loop *de novo* modelling results on TyrRS mutants with PDB ID 1ZH0, 1ZH6, 2AG6 and 3D6U, respectively. (A) Box plot of rmsd_to_crystal for the top 10 models ranked by total_energy_score on 4 TyrRS mutants. (B) Box plot of total_score rank for the top 10 models ranked by rmsd_to_crystal on 4 TyrRS mutants. (C, D, E, F) Structure superimposition of crystal structure (in green), original Rosetta model using WT crystal structure as template (in blue), remodeled model with the lowest RMSD to crystal structure within the 1000 models (in magenta) and remodeled model with the lowest energy within the 1000 models (in yellow) on 4 TyrRS mutants.

### 2.3 Machine learning model and performance analysis

In the previous part, we have modeled 6864 UAA-TyrRS mutant pairs using Rosetta and found that the energy function is not accurate enough to discriminate the true UAA binder from the false. To further improve the energy function, we use machine learning to train a model to learn important energy term and protein-ligand interactions that contribute most to the binding energy. The flowchart of the machine learning model and application is shown in Scheme 2. Next each part of the flowchart will be described.

**Scheme 2.**
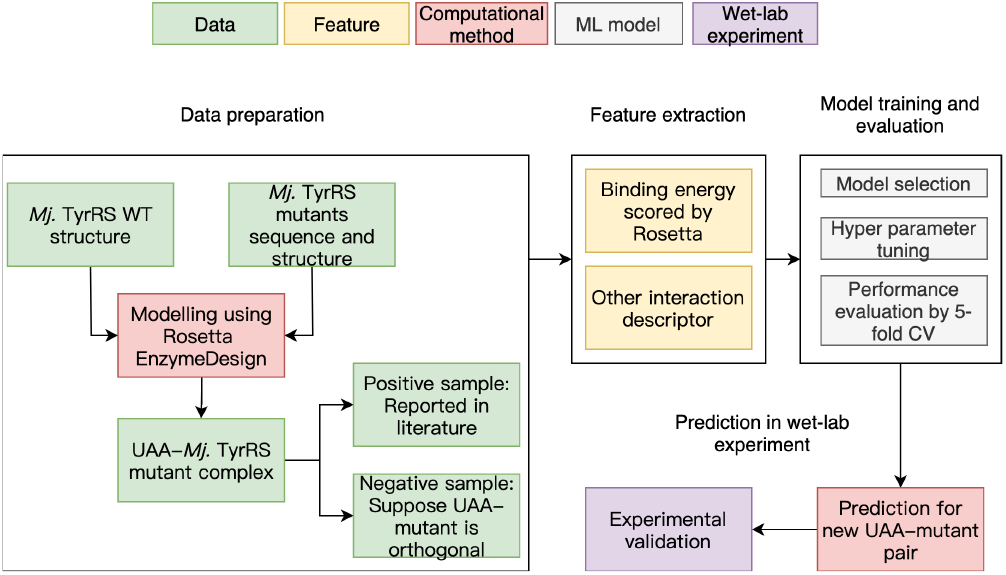
Flowchart of the data preparation, ML model training, prediction and application workflow. Figure legend is on the top.

#### 2.3.1 Data preparation

We collected the chemical structure of 52 UAAs, sequence and X-ray crystal structure of 132 TyrRS mutants from previous literature. Each UAA is paired to each TyrRS mutant to generate data set for ML model training. There are 6864 UAA-TyrRS mutant pairs and complex model is built in the previous section. For the specific UAA-mutant pair, if the UAA can be recognized by the mutant, this pair is labeled as positive and all the remaining pairs are taken as negative sample under the assumption that TyrRS mutant has high specificity and orthogonality for different UAA substrate. In the final dataset, we have 132 positive samples and 6732 negative samples. The positive to negative ratio is 1:51.

#### 2.3.2 Feature extraction

The UAA-TyrRS mutant pair can be described from 3 aspects: UAA ligand, mutant protein and UAA-mutant complex. The presentation and feature vectorization method are listed in Table 3. The feature is extracted from the chemical structure of UAA, sequence of TyrRS mutant and 3D structure of UAA-mutant complex model built using Rosetta. The dimension of all features is 2644.

**Table 3.** Feature representation method for UAA small molecule ligand and TyrRS protein receptor used in this study

| Method | Description | Dimension | Reference |
| --- | --- | --- | --- |
| PROFEAT-ligand | Small molecule 1D and 2D descriptors. | 406 | 41 |
| PROFEAT-receptor | Protein sequence descriptors using amino acid biophysical properties and compositions. | 1437 | 41 |
| Rosetta energy score and decomposition | Rosetta interface_E total score and energy term (fa_atr, fa_rep, fa_sol, fa_elec, fa_pair, hbond_sr_bb, hbond_lr_bb, hbond_bb_sc, hbond_sc) decomposed into each binding pocket residue. | 6+540 | 22, 42 |
| dpocket | Dpocket (describing pocket) extracts several descriptors using atom, amino acid, alpha sphere and volume information from the ligand binding pocket. | 35 | 43 |
| rfscore | A machine learning-based score function for protein-ligand complex. | 216 | 44 |

#### 2.3.3 ML Model training

10% of data is used as test set (687 samples). The remaining 90% data (6177 samples) is used for ML model training, which is randomly splitted into training set and validation set with a ratio of 4:1 for 5-fold cross validation (CV). Feature engineering, model selection and hyper parameter tuning are carried out through pycaret^40^, a low-code machine learning library. 15 machine learning algorithms are used to train the model: Light Gradient Boosting Machine (lightGBM), CatBoost Classifier, Extra Trees Classifier, Extreme Gradient Boosting, Logistic Regression, Ridge Classifier, Random Forest Classifier, Quadratic Discriminant Analysis, K Neighbors Classifier, Ada Boost Classifier, Gradient Boosting Classifier, Linear Discriminant Analysis, Decision Tree Classifier, SVM - Linear Kernel, Naive Bayes Classifier. Different metrics are used to evaluate the binary classification model performance: accuracy, AUC, recall, precision and F1.

#### 2.3.4 ML Model evaluation and explanation

Different feature combinations are explored in the model training process. The feature_type_index, values of best metrics and corresponding model name having the best performance are listed in Table 4. For different metrics, the algorithm having best performance is different. In general, the performance of model using all features is better than using single type of feature (Figure 8). When using the UAA dpocket descriptor only, Quadratic Discriminant Analysis and Extra Trees Classifier models have best recall and precision, respectively, while their F1 is not the highest. LightGBM model has the best accuracy, AUC and precision in some feature combinations.

**Table 4.** Comparison of 5-fold CV performance on different ML models and different performance metrics for UAA specificity prediction

| Features and combination | Feature_type_index | Model_with_best_accuracy | Best_accuracy | Model_with_best_AUC | Best_AUC | Model_with_best_recall | Best_recall | Model_with_best_precision | Best_precision | Model_with_best_F1 | Best_F1 |
| --- | --- | --- | --- | --- | --- | --- | --- | --- | --- | --- | --- |
| Rosetta_6_score_terms | 1 | SVM - Linear Kernel | 0.979 | CatBoost Classifier | 0.7369 | Decision Tree Classifier | 0.0615 | Naive Bayes | 0.176 | Quadratic Discriminant Analysis | 0.0777 |
| Rosetta_energy_decomposed | 2 | Extra Trees Classifier | 0.9796 | Light Gradient Boosting Machine | 0.7874 | Naive Bayes | 0.8077 | Random Forest Classifier | 0.4000 | Decision Tree Classifier | 0.1465 |
| UAA_pocket_descriptotr | 3 | Extra Trees Classifier | **0.9799** | CatBoost Classifier | 0.8042 | Quadratic Discriminant Analysis | 1.0000 | Extra Trees Classifier | 0.7167 | Extra Trees Classifier | 0.1612 |
| Rosetta_6_score_terms+energy_decomposed | 4 | Light Gradient Boosting Machine | 0.9801 | CatBoost Classifier | 0.8193 | Linear Discriminant Analysis | 0.1846 | K Neighbors Classifier | 0.6500 | Linear Discriminant Analysis | 0.1836 |
| All_features_above | 5 | Light Gradient Boosting Machine | **0.9794** | CatBoost Classifier | 0.8222 | Naive Bayes | 0.4769 | Light Gradient Boosting Machine | 0.4000 | Linear Discriminant Analysis | 0.1654 |

**Figure 8.**
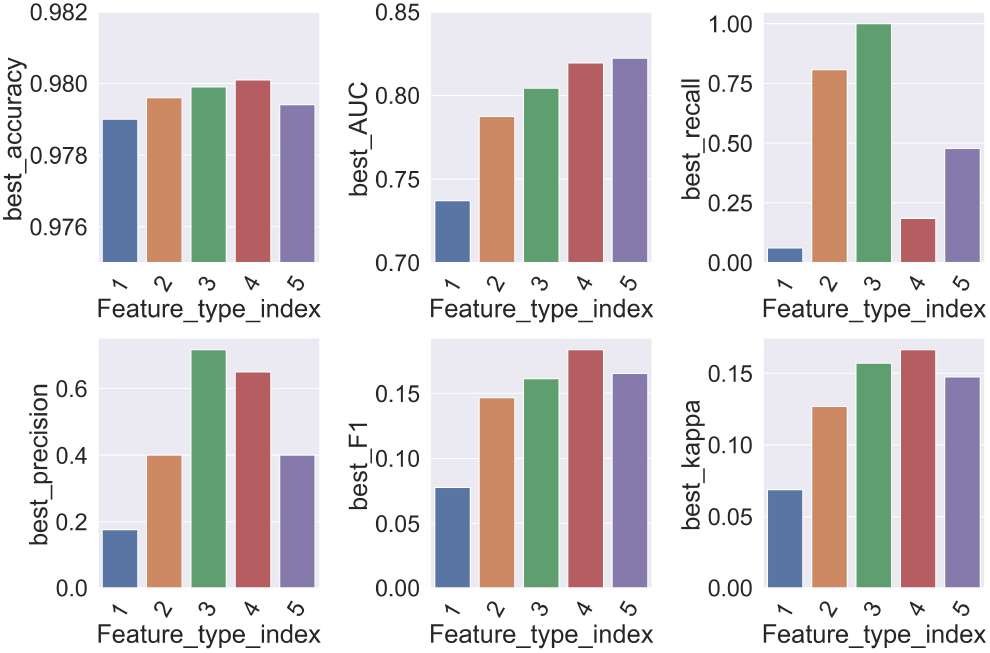
Barplot of the best model performance on different feature combinations and different metrics. Feature_type_index is the same as that in Table 4.

We choose lightGBM^45^ model for further analysis since it’s a tree-based ensemble model which can avoid overfitting and has been widely used in other machine learning model applications. Feature importance of the model can be easily explained. The performance of lightGBM model on 5-fold CV and test set is compared (Table 5). We can see the metrics are better on test set than that on 5-fold CV.

**Table 5.** Performance of lightGBM model on 5-fold cross validation set and test set using all features (feature_type_index 5).

| lightGBM | Accuracy | AUC | Recall | Precision | F1 |
| --- | --- | --- | --- | --- | --- |
| 5-fold CV | 0.9794 | 0.7593 | 0.0385 | 0.35 | 0.0686 |
| test set | 0.9796 | 0.8549 | 0.0667 | 1 | 0.125 |

SHAP method^46^ is used to calculate the feature importance and its contribution to the binary class label (Figure 9). Residue_70_fa_sol and residue_34_fa_sol are considered as the most important features by the lightGBM model which indicates that the solvation energy of residue 34 and 70 may be important. Rosetta_interface_E and residue_158_fa_elec are also important which is in accordance with our domain knowledge that score function is useful to discriminate the positive sample to some extent.

**Figure 9.**
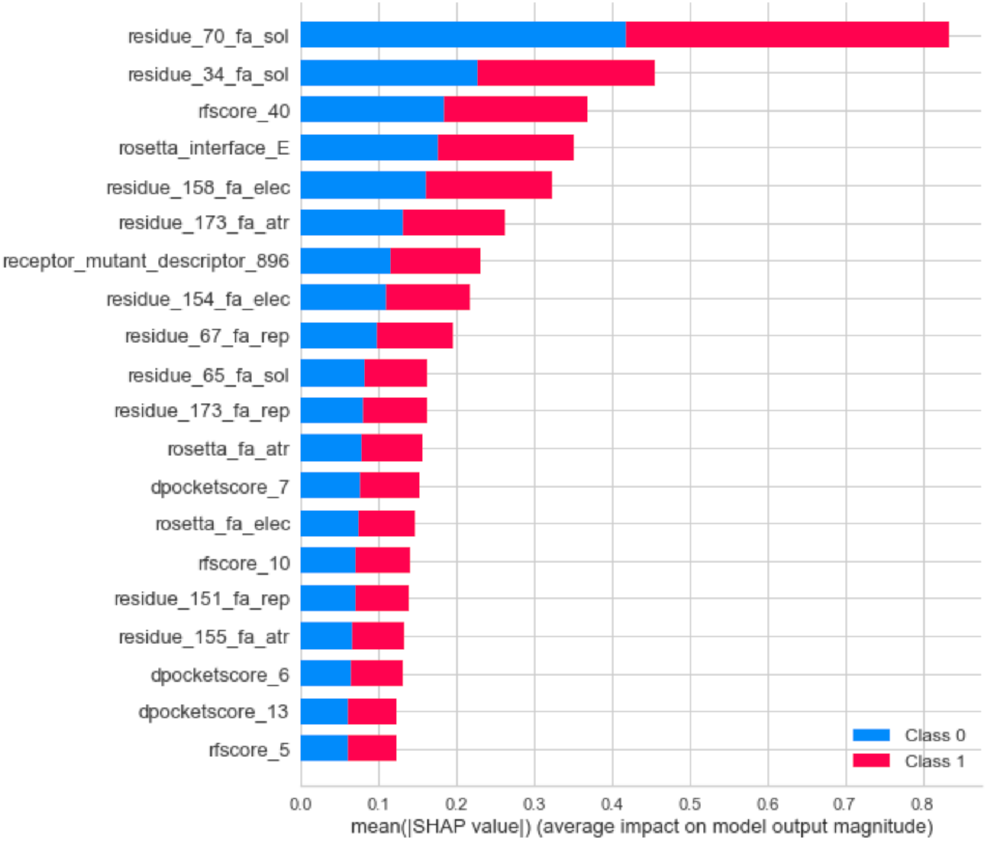
SHAP values of the important features for the lightGBM model. The average SHAP value of specific feature in all samples are shown. The impact on negative label is in blue and the impact on positive label is in red.

To compare the prediction accuracy between lightGBM model and Rosetta score, ROC and PR curve on test set (Figure 10A and 10B) are plotted for both 2 methods. Better ROC and PR curve are observed for lightGBM ML model. AUC of lightGBM model is 0.84, higher than 0.77 of Rosetta score. As a simulation test for real wet lab experiment, we calculate when the number of mutants k for web-lab exp is given and fixed, how many true positive samples exist in the k mutants (Figure 10C and 10D). For example, if we want to test 50 mutants in wet lab experiment out of 687 mutants, there will be 1, 2 and 11 true positive mutants for random sample, Rosetta score prediction and lightGBM model prediction, respectively. The success rate is 2%, 4% and 22%, respectively, which means Rosetta score prediction has 2-fold elevation on enrichment ratio of true positive mutants, while lightGBM model has 11-fold elevation. The TyrRS mutant screening will significantly benefit from the improvement of prediction accuracy using machine learning model.

**Figure 10.**
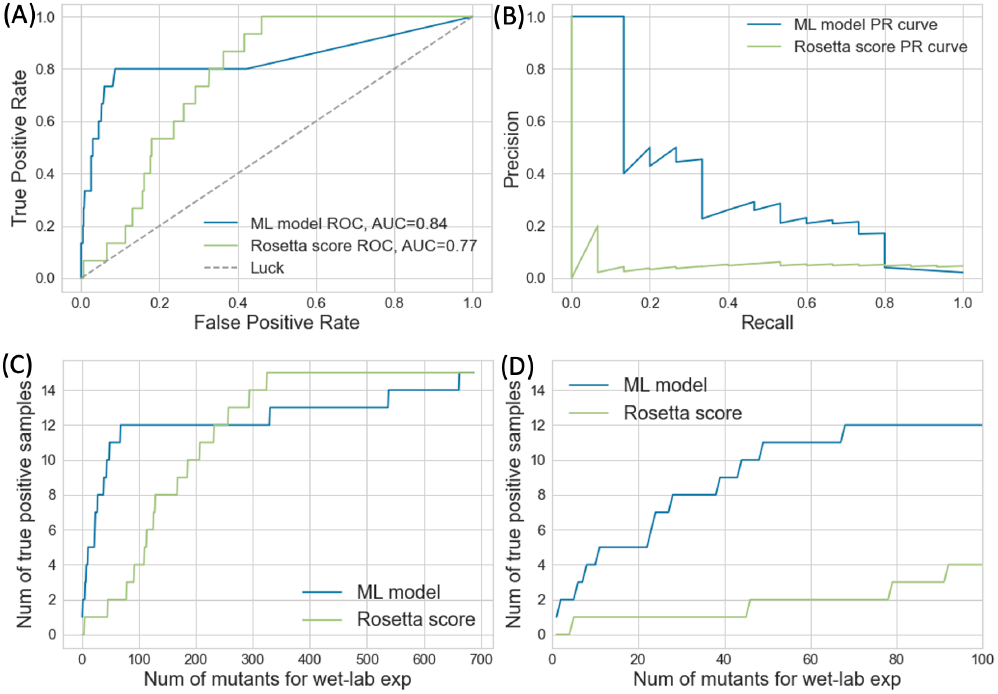
(A) Receiver operating characteristic curve, (B) Precision-recall curve and (C, D) Number of true positive samples vs number of mutants selected for wet-lab experiment of lightGBM model and Rosetta interface_E score only prediction on test set. The scale of x axis is limited to 0-100 in figure D.

## 3 Discussion

### 3.1 Significance of the work for genetic code expansion and computational protein design

In the field of computational protein design, it is a great challenge to introduce mutations into proteins to change the substrate specificity for different ligands in a particular ligand-receptor complex system. There have been many reports, such as using Rosetta^47^ and OSPREY^48^ to design proteins to switch the substrate and accommodate specific ligands. As a model system, *Mj.* TyrRS has be mutated and designed to bind UAA with different chemical structures as reported before^49^. This system can be used as a benchmark for substrate-specific protein design, as a large number of mutants have been reported. Comprehensive study of this system will be of great significance to the field of protein design.

This work focuses on UAA-*Mj.* TyrRS complex system which has been widely used in genetic code expansion research field. Through the efforts of many research groups in the past over 15 years, a large number of UAA and its corresponding TyrRS mutants has been identified after spending a lot of human and material resources, but so far, there is no systematic study to integrate these data for the development and test of UAA virtual screening method. This paper summarizes all the reported UAA and *Mj.* TyrRS mutants, systematically analyzes them, and attempts to establish a relatively accurate model to predict the UAA-specific recognition of different mutants, establishing the foundation for large-scale, high-throughput virtual screening.

The existing methods for predicting protein ligand interaction are probably not suitable for this system. The reason is that the adaptability and accuracy of score functions for different ligand-receptor systems are different. For example, the importance of hydrogen bond in one system may be higher than another. Method developed for affinity prediction of drug-target protein interaction cannot be applied on the *Mj.* TyrRS system without modification. In this paper, we calibrate the Rosetta score function for this specific system to get better prediction performance. Besides Rosetta molecular modeling, in the future other methods such as molecular docking, MM-PBSA^50^, TI^51^, and FEP^52^ can also be integrated with ML in the similar way to give more accurate prediction results.

### 3.2 Previous work for TyrRS mutant substrate specificity prediction and computational library design

Several methods aiming to accelerate the screening of aaRS mutants or design more focusing library using computational chemistry method such as molecular modeling and MM-PBSA^53–55^ have been reported. These results show that molecular modeling can be used to predict TyrRS substrate specificity, but the amount of UAA and TyrRS mutants used in previous studies is too small to give convincing conclusion. In this work, we considered all UAA and TyrRS mutants reported before. With thousands of UAA-mutant pairs as training and validation data, a more solid conclusion can be made that molecular modeling integrated with ML can be useful in the virtual screening of TyrRS mutant.

### 3.3 Discussion on the prediction result

Though the backbone disruption of helix 158-163 can’t be precisely modeled, we think its influence on the prediction accuracy may not be large because there are only 58 D158G mutants and 13 D158P mutants with a fraction of 42% in total 171 mutants (Supplementary file S1). Besides, descriptors extracted from protein sequence is not affected by the backbone disruption of structure. Knowledge and information learned from mutant sequence by ML model could correct the error in the homology model and increase the prediction accuracy.

### 3.4 Limitation of the work

#### Limitation of the current molecular modeling method

Firstly, for the homology modeling of *Mj.* TyrRS mutants, the position of amino acid side chain and rotamer in the binding pocket can be recovered well, but for those with large backbone changes, it is hard to model the backbone conformation aligned well to crystal structure. Though the close conformation can be sampled by Rosetta loop remodeling, the score function is not precise enough to pick it out. The inaccuracy of structure prediction decreases the prediction accuracy of UAA-specific mutant. Secondly, there are water molecules in the UAA binding pocket of *Mj.* TyrRS. The influence of water is not considered by current molecular modeling method.

#### Limitation of the machine learning method

The generalization ability of ML model may be low because the number of reported UAA and mutant pair is small compared to the huge chemical space of UAA and sequence space of *Mj.* TyrRS mutants. Currently it is difficult to cover the diversified chemical structure and protein sequence. One of the solutions is deep mutational scanning^56^, which uses next generation sequencing to get phenotype of over 1 million mutants in a single experiment and generates enough data for machine learning and deep learning model training and prediction. Still wet-lab experiments are needed to validate the model for practical usage.

## 4 Conclusion

To get further knowledge of the *Mj.* TyrRS structure and accelerate the time-costly screening process of mutant for specific UAA, we collected all the UAAs and *Mj.* TyrRS reported before, analyzed the structure and sequence difference between the mutants and found that some mutants have alternative backbone conformation on alpha helix residue 158-163 with D158G/P mutation, which makes accurate mutant modelling more difficult. Rosetta modeling and machine learning are integrated to give more accurate prediction results for mutant selectivity towards different UAAs. Different feature combinations and machine learning algorithms are tested for higher model performance. Lightgbm model is chosen and the feature importance and contribution to the binary class label is calculated to explain the knowledge learned by the model. After the calibration of Rosetta score function using lightGBM model, the enrichment ratio of target mutant is elevated by 11-fold compared with random mutation. Wet-lab experiment is in progress to validate the model. We anticipate that this proof-of-concept workflow will be of great help in the screening of *Mj*. TyrRS and protein design field.

## 5 Methods

### 5.1 Molecular modelling using Rosetta

#### Preparation of the UAA ligand

Preparation of the UAA ligand was performed as described in the Rosetta tutorial for ligand preparation^57^. Briefly speaking, the chemical structure of UAA is drawn using Marvin JS^58^ and converted to smiles format. Cheminformatics tool RDKit^59^ is used to generate low-energy conformations and perform energy minimization based on the smiles of UAA. The ligand parameter file used in Rosetta is made by molfile_to_params.py^57^.

#### UAA-*Mj.* TyrRS mutant complex model building

The UAA-*Mj.* TyrRS mutant complex model is built using Rosetta Enzyme design^22^ application with default parameters as described in Rosetta tutorial^60^. Briefly speaking, the amino acid mutation in the mutants is written in resfile and passed into the application along with the UAA ligand parameter file. Catalytic residues of the aaRS formatting hydrogen bond with UAA is fixed using constraint file. For each complex, nstruct = 10 is used and the structure with the lowest total score is kept for further analysis.

#### De novo loop modeling using KIC method

The residues 158-163 segment of *Mj.* TyrRS is *de novo* modeled using loop modeling KIC method as described in Rosetta tutorial^61^ with default parameters. 1000 models are generated for each input structure. The RMSD between modeled structure and crystal structure is calculated by in-house script written using biopython^62^.

### 5.2 Machine learning

#### Descriptor extraction

PROFEAT, dpocket and rfscore descriptors of UAA ligand, *Mj.* TyrRS mutant protein and UAA-*Mj.* TyrRS mutant complex is calculated on http://www.descriptordb.com/ with default parameters.

#### Feature engineering

Feature engineering, model selection and hyperparameter optimization are performed by PyCaret^40^. 90% of the full data is used in 5-fold CV and the rest 10% of data is used as test set. The feature space is transformed using ‘zscore’ method. ‘ignore_low_variance’ option is set to True and all categorical features with statistically insignificant variances are removed from the dataset. Feature selection is used and ‘feature_selection_threshold’ is set to 0.8.

## Data availability

All datasets and parameters generated or analyzed during the study are available from the corresponding author on reasonable request.

## Author contributions

Yingfei Sun conceived the study and edited the paper. Bingya Duan conducted the experiments and analyzed the results. Bingya Duan and Yingfei Sun co-wrote the manuscript. All authors reviewed the manuscript.

## References

1. J. W. Chin, Expanding and reprogramming the genetic code of cells and animals. Annu Rev Biochem 83, 379–408 (2014).

2. F. Yang et al., Phospho-selective mechanisms of arrestin conformations and functions revealed by unnatural amino acid incorporation and (19)F-NMR. Nat Commun 6, 8202 (2015).

3. F. Li et al., A genetically encoded 19F NMR probe for tyrosine phosphorylation. Angew Chem Int Ed Engl 52, 3958–3962 (2013).

4. K. Yokoyama, U. Uhlin, J. Stubbe, Site-Specific Incorporation of 3-Nitrotyrosine as a Probe of pK(a) Perturbation of Redox-Active Tyrosines in Ribonucleotide Reductase. J Am Chem Soc 132, 8385–8397 (2010).

5. I. N. Ugwumba et al., Improving a Natural Enzyme Activity through Incorporation of Unnatural Amino Acids. J Am Chem Soc 133, 326–333 (2011).

6. I. Drienovska, G. Roelfes, Expanding the enzyme universe with genetically encoded unnatural amino acids. Nat Catal 3, 193–202 (2020).

7. X. H. Liu et al., A genetically encoded photosensitizer protein facilitates the rational design of a miniature photocatalytic CO2-reducing enzyme. Nat Chem 10, 1201–1206 (2018).

8. I. Drienovska et al., Design of an enantioselective artificial metallo-hydratase enzyme containing an unnatural metal-binding amino acid. Chem Sci 8, 7228–7235 (2017).

9. Q. Li et al., Developing Covalent Protein Drugs via Proximity-Enabled Reactive Therapeutics. Cell, (2020).

10. V. K. Vyas, R. D. Ukawala, M. Ghate, C. Chintha, Homology Modeling a Fast Tool for Drug Discovery: Current Perspectives. Indian J Pharm Sci 74, 1–17 (2012).

11. A. W. Senior et al., Improved protein structure prediction using potentials from deep learning. Nature 577, 706–+ (2020).

12. X. Y. Meng, H. X. Zhang, M. Mezei, M. Cui, Molecular Docking: A Powerful Approach for Structure-Based Drug Discovery. Curr Comput-Aid Drug 7, 146–157 (2011).

13. B. Kuhlman, P. Bradley, Advances in protein structure prediction and design. Nat Rev Mol Cell Bio 20, 681–697 (2019).

14. R. F. Li et al., Computational redesign of enzymes for regio- and enantioselective hydroamination. Nat Chem Biol 14, 664–+ (2018).

15. A. Zhavoronkov et al., Deep learning enables rapid identification of potent DDR1 kinase inhibitors. Nat Biotechnol 37, 1038–+ (2019).

16. Z. Wu, S. B. J. Kan, R. D. Lewis, B. J. Wittmann, F. H. Arnold, Machine learning-assisted directed protein evolution with combinatorial libraries (vol 116, pg 8852, 2019). P Natl Acad Sci USA 117, 788–789 (2020).

17. J. A. Ruffolo, C. Guerra, S. P. Mahajan, J. Sulam, J. J. J. b. Gray, Geometric Potentials from Deep Learning Improve Prediction of CDR H3 Loop Structures. (2020).

18. J. Graves et al., A Review of Deep Learning Methods for Antibodies. Antibodies (Basel) 9, (2020).

19. C. Wang, Y. Zhang, Improving scoring-docking-screening powers of protein-ligand scoring functions using random forest. J Comput Chem 38, 169–177 (2017).

20. J. X. Wang, H. L. Cao, J. Z. H. Zhang, Y. F. Qi, Computational Protein Design with Deep Learning Neural Networks. Sci Rep-Uk 8, (2018).

21. H. Cao, J. Wang, L. He, Y. Qi, J. Z. Zhang, DeepDDG: Predicting the Stability Change of Protein Point Mutations Using Neural Networks. J Chem Inf Model 59, 1508–1514 (2019).

22. F. Richter, A. Leaver-Fay, S. D. Khare, S. Bjelic, D. Baker, De novo enzyme design using Rosetta3. PLoS One 6, e19230 (2011).

23. Schrodinger, LLC. (2015).

24. T. Kobayashi et al., Structural basis for orthogonal tRNA specificities of tyrosyl-tRNA synthetases for genetic code expansion. Nat Struct Biol 10, 425–432 (2003).

25. Y. Zhang, L. Wang, P. G. Schultz, I. A. Wilson, Crystal structures of apo wild-type M. jannaschii tyrosyl-tRNA synthetase (TyrRS) and an engineered TyrRS specific for O-methyl-L-tyrosine. Protein Sci 14, 1340–1349 (2005).

26. J. M. Turner, J. Graziano, G. Spraggon, P. G. Schultz, Structural plasticity of an aminoacyl-tRNA synthetase active site. Proc Natl Acad Sci U S A 103, 6483–6488 (2006).

27. W. Liu, L. Alfonta, A. V. Mack, P. G. Schultz, Structural basis for the recognition of para-benzoyl-L-phenylalanine by evolved aminoacyl-tRNA synthetases. Angew Chem Int Ed Engl 46, 6073–6075 (2007).

28. J. M. Turner, J. Graziano, G. Spraggon, P. G. Schultz, Structural characterization of a p-acetylphenylalanyl aminoacyl-tRNA synthetase. J Am Chem Soc 127, 14976–14977 (2005).

29. D. D. Young et al., An evolved aminoacyl-tRNA synthetase with atypical polysubstrate specificity. Biochemistry 50, 1894–1900 (2011).

30. E. M. Tippmann, W. Liu, D. Summerer, A. V. Mack, P. G. Schultz, A genetically encoded diazirine photocrosslinker in Escherichia coli. Chembiochem 8, 2210–2214 (2007).

31. J. Xie, W. Liu, P. G. Schultz, A genetically encoded bidentate, metal-binding amino acid. Angew Chem Int Ed Engl 46, 9239–9242 (2007).

32. R. B. Cooley et al., Structural basis of improved second-generation 3-nitro-tyrosine tRNA synthetases. Biochemistry 53, 1916–1924 (2014).

33. J. Wang et al., A biosynthetic route to photoclick chemistry on proteins. J Am Chem Soc 132, 14812–14818 (2010).

34. X. Liu et al., Significant expansion of fluorescent protein sensing ability through the genetic incorporation of superior photo-induced electron-transfer quenchers. J Am Chem Soc 136, 13094–13097 (2014).

35. R. B. Cooley, P. A. Karplus, R. A. Mehl, Gleaning unexpected fruits from hard-won synthetases: probing principles of permissivity in non-canonical amino acid-tRNA synthetases. Chembiochem 15, 1810–1819 (2014).

36. X. Luo et al., Genetically encoding phosphotyrosine and its nonhydrolyzable analog in bacteria. Nat Chem Biol 13, 845–849 (2017).

37. S. Kawashima et al., AAindex: amino acid index database, progress report 2008. Nucleic Acids Res 36, D202–205 (2008).

38. F. Pedregosa et al., Scikit-learn: Machine Learning in Python. 12, 2825–2830 (2011).

39. A. Stein, T. Kortemme, Improvements to robotics-inspired conformational sampling in rosetta. PLoS One 8, e63090 (2013).

40. M. Ali, PyCaret: An open source, low-code machine learning library in Python. (2020).

41. Z. R. Li et al., PROFEAT: a web server for computing structural and physicochemical features of proteins and peptides from amino acid sequence. Nucleic Acids Res 34, W32–37 (2006).

42. R. F. Alford et al., The Rosetta All-Atom Energy Function for Macromolecular Modeling and Design. J Chem Theory Comput 13, 3031–3048 (2017).

43. P. Schmidtke, V. Le Guilloux, J. Maupetit, P. Tuffery, fpocket: online tools for protein ensemble pocket detection and tracking. Nucleic Acids Res 38, W582–589 (2010).

44. P. J. Ballester, J. B. Mitchell, A machine learning approach to predicting protein-ligand binding affinity with applications to molecular docking. Bioinformatics 26, 1169–1175 (2010).

45. G. Ke et al., in neural information processing systems. (2017), pp. 3149–3157.

46. S. Lundberg et al., From local explanations to global understanding with explainable AI for trees. 2, 56–67 (2020).

47. R. Moretti, B. J. Bender, B. Allison, J. Meiler, Rosetta and the Design of Ligand Binding Sites. Methods Mol Biol 1414, 47–62 (2016).

48. M. A. Hallen et al., OSPREY 3.0: Open-source protein redesign for you, with powerful new features. J Comput Chem 39, 2494–2507 (2018).

49. J. W. Chin, Expanding and reprogramming the genetic code. Nature 550, 53–60 (2017).

50. S. Genheden, U. Ryde, The MM/PBSA and MM/GBSA methods to estimate ligand-binding affinities. Expert Opin Drug Dis 10, 449–461 (2015).

51. T. S. Lee, Y. Hu, B. Sherborne, Z. Guo, D. M. York, Toward Fast and Accurate Binding Affinity Prediction with pmemdGTI: An Efficient Implementation of GPU-Accelerated Thermodynamic Integration. J Chem Theory Comput 13, 3077–3084 (2017).

52. Z. Cournia, B. Allen, W. Sherman, Relative Binding Free Energy Calculations in Drug Discovery: Recent Advances and Practical Considerations. Journal of Chemical Information and Modeling 57, 2911–2937 (2017).

53. W. Ren, T. M. Truong, H. W. Ai, Study of the Binding Energies between Unnatural Amino Acids and Engineered Orthogonal Tyrosyl-tRNA Synthetases. Sci Rep-Uk 5, (2015).

54. V. Opuu, G. Nigro, E. Schmitt, Y. Mechulam, T. Simonson, Adaptive landscape flattening allows the design of both enzyme:substrate binding and catalytic power. 771824 (2019).

55. T. Baumann et al., Computational Aminoacyl-tRNA Synthetase Library Design for Photocaged Tyrosine. Int J Mol Sci 20, (2019).

56. D. M. Fowler, S. Fields, Deep mutational scanning: a new style of protein science. Nat Methods 11, 801–807 (2014).

57. P. Hosseinzadeh, Preparing Ligands, https://www.rosettacommons.org/demos/latest/tutorials/prepare_ligand/prepare_ligand_tutorial. (2016).

58. https://marvinjs-demo.chemaxon.com/latest/.

59. G. Landrum, RDKit: Open-source cheminformatics, http://www.rdkit.org. (2020).

60. F. Richter, Enzyme design application, https://www.rosettacommons.org/docs/latest/application_documentation/design/enzyme-design. (2010).

61. A. Stein, Next-generation kinematic loop modeling and torsion-restricted sampling, https://www.rosettacommons.org/docs/latest/application_documentation/structure_prediction/loop_modeling/next-generation-KIC. (2015).

62. P. J. Cock et al., Biopython: freely available Python tools for computational molecular biology and bioinformatics. Bioinformatics 25, 1422–1423 (2009).

